# Centrosome-assisted assembly of the Balbiani body

**DOI:** 10.1101/2025.02.11.637656

**Authors:** Divyanshi, Meining Li, Elvan Böke, Jing Yang

## Abstract

The Balbiani body (Bb), which was discovered about 170 years ago, is a membraneless organelle in the oocyte in most species. In organisms like *Xenopus* and Zebrafish, Bb accumulates mitochondria, endoplasmic reticulum (ER), and germline determinants and regulates the proper localization of germline determinants. The Bb forms around the centrosome in the oocyte during early oogenesis. The mechanism behind its assembly has gained attention only very recently. Here, we report that overexpression of the germ plasm matrix protein Xvelo leads to the formation of a ‘Bb-like’ structure in somatic cells. The ‘Bb-like’ structure assembles around the centrosome and selectively recruits mitochondria, ER, and germline determinants. Taking advantage of this system, we investigated the roles of centrosome components on the assembly of Xvelo. Our results reveal that multiple components of the centrosome, including Sas6, Cenexin, and DZIP1, interact with Xvelo and promote its assembly, with Sas6 exhibiting the most prominent activity. Importantly, knocking down Sas6, Cenexin, and DZIP1 individually or in combination resulted in reduced Xvelo aggregates. Taken together, our work suggests that the centrosome may function as a nucleation center to promote the initiation of Xvelo assembly, resulting in the formation of the Bb around the centrosome.

## Introduction

Recent works have highlighted the fundamental roles of membraneless biomolecular condensates in a wide variety of biological processes^1–3^. By enriching biomolecules in specific subcellular compartments, membraneless condensates can either facilitate biochemical reactions inside the cell or regulate the half-life of their cargoes. Increasing evidence has demonstrated that under physiological conditions, membraneless condensates are involved in gene regulation at transcriptional, translational, and post-translational levels^4–7^. However, formation of pathological condensates can be toxic to the cell, leading to devastating conditions such as neurodegenerative diseases in humans^8,9^.

Membraneless condensates are fundamental for germline development^10,11^. Several condensates unique to the germline have been identified across different species like the P granules or germline P bodies in the worm^12^, polar granules in the fly^10,13^, L body in the frog^14,15^ and male (XY) sex body in mammals^16–18^. Apart from these dynamic gel/liquid-like condensates, stable amyloid-like protein assemblies also exist in the germline. Unlike pathological amyloids that accumulate under disease conditions^8,9^, these amyloid-like aggregates form under physiological conditions. In yeast, dynamic regulation of such condensates is essential for gametogenesis^19,20^. The Balbiani body (Bb), an amyloid-like condensate, first discovered more than a hundred years ago^21,22^, has been observed in the oocytes in many species^10,23–25^.

The Bb has been studied by several generations of researchers^22,24,26,27^. Studies in the 1800s first identified it as granulofibrillar material in the oocytes of spiders^21^. It was later observed in *Drosophila*, *Xenopus*, zebrafish, mice, and human oocytes^28–33^. The conservation of this structure in oocytes through the course of evolution hints at its potential importance. Although the precise arrangement varies across species, the Bb houses abundant mitochondria, endoplasmic reticulum (ER), Golgi apparatus, and/or specific set of RNAs^22,28,29,34,35^. It may play a role in ensuring the inheritance of healthy organelles by the next generation, which could explain the need for its conservation across species^36–38^. In *Xenopus* and zebrafish, the Bb accumulates the RNAs and proteins needed to specify future primordial germ cells^33,39–42^. *Xenopus* Velo (Xvelo) and zebrafish Buckyball (Buc) proteins form the matrix for the Bb in these species^30,33,39,42,43^. In zebrafish buc null mutants, Bb cannot assemble in the oocyte. The mitochondria, ER, and germline determinants are dispersed in the entire cytoplasm. Likely due to the defective animal-vegetal axis, embryos derived from *buc^-/-^* oocytes cannot form blastodisc and die shortly after fertilization^30,33,44^.

The Xvelo protein contains an N-terminal Prion-like domain (PLD) and a C-terminal intrinsically disordered region (IDR)^39,49^. In general, proteins with IDRs or PLD have a high propensity to take part in the process of phase separation^1,39,45–49^. It has been reported that the N-terminus PLD of Xvelo, which confers Xvelo the ability to self-assemble, is required for its localization to the Bb in *Xenopus* oocytes. The efficient recruitment of RNA and organelles into the Bb, however, also requires the C-terminus of the protein^39^. While the Xvelo/Buc protein can self-assemble into an amyloid-like network in vitro^39^ and is required for Bb formation in the oocyte^30,33,44^, it is currently unclear whether Xvelo/Buc alone can assemble into a Bb in the absence of other oocyte specific factors. Although Bb assembly has started to gain attention recently^50^, the mechanism responsible for the initiation of this assembly during early oogenesis remains largely unknown.

In *Xenopus* and zebrafish, the Bb assembles around the centrosome in stage I oocytes^51,52^. The centrosome is a microtubule organizing center (MTOC) in the cell and consists of a pair of centrioles surrounded by the pericentriolar matrix (PCM)^53,54^. In most species, centrioles are eliminated from the oocyte during later stages of oogenesis and are reintroduced by the sperm upon fertilization^55–58^. The centrosome is composed of hundreds of proteins. Many of these proteins have been studied during early development. For instance, Polo-like kinase 4 (PLK4), which can partake in self-assembly via phase separation^59,60^, was shown to play an important role in the regulation of centriole duplication during early zebrafish embryogenesis^61^. The outer dense fiber (ODF2) isoform 9, also called Cenexin, was recently reported to play a role in maintaining the integrity of the PCM in zebrafish embryos^62^. Perturbations to some centrosome proteins have also been shown to cause defects in the germline. Zebrafish embryos mutant for Spindle assembly defective protein 6 (Sas6), which acts as the scaffold for the centrioles^63–65^, showed defective germ plasm recruitment to the cleavage furrows in early embryos^66^. Dzip1 is another component of the appendages at mother centrioles with documented roles in both germline and somatic development. In somatic tissues, Dzip1 regulates ciliogenesis and is essential for the Hedgehog (Hh) signaling in early embryos^67–74^. Dzip1 is dynamically expressed in vertebrate germline. It localizes to the Bb in *Xenopus* and zebrafish oocytes. Knockdown of Dzip1 impairs the development of *Xenopus* primordial germ cells^75^. So far, whether Dzip1 or any other centriolar proteins play a role in Bb assembly remains unknown.

In this study, we report that Xvelo possesses the ability to assemble into a Bb-like structure around the centrosome in mammalian somatic cells. These Xvelo assemblies accumulate mitochondria and ER and can selectively enrich co-expressed germline proteins. Centrosome components such as Sas6, Cenexin, and DZIP1 interact with Xvelo and enhance the assembly of Xvelo. Importantly, the knockdown of these centriole components reduced the Xvelo assembly. Our results support the idea that assembly of Xvelo, aided by interactions with centrosome components, could initiate Bb formation around the centrosome, providing novel mechanistic insight into the initial steps of Bb formation.

## Results

### Overexpression of Xvelo induces a ‘Bb-like’ structure in mammalian somatic cells

The Bb assembles around the centrosome in early-stage oocytes^51,52^. In *Xenopus*, Xvelo protein forms the stable matrix of the Bb and was reported to show minimal recovery of fluorescence post-photobleaching^39^. We found that when GFP-Xvelo was expressed in HEK293T cells, Xvelo in some cells assembled into aggregates (SFig1A). The assembly of Xvelo aggregates is dose-dependent. At the condition used for the majority of experiments in this study (0.5µg plasmid per transfection), around 10% of Xvelo-expressing cells formed Xvelo aggregates and the rest showed a smear Xvelo expression pattern (SFig1A-B). To assess Xvelo assembly more quantitatively, we performed a fractionation experiment in which Xvelo-expressing cells were lysed with the NP40-containing lysis buffer, followed by centrifugation to separate the soluble (smear Xvelo) and insoluble (Xvelo aggregates) protein (SFig1C). The expression level of insoluble Xvelo was undetectable in cells transfected with 0.25 µg Xvelo expression construct. In cells transfected with 1.5 µg Xvelo plasmid, however, we found around half of Xvelo was in the insoluble form (SFig1C). In these transient transfection experiments, Xvelo assembles over time, before eventually being degraded (SFig1D). The Xvelo assemblies were resistant to digitonin extraction. This is in stark contrast to the GFP protein in the cytoplasm, which was extracted completely upon digitonin permeabilization of the cell (SFig1E).

Strikingly, Xvelo often assembles in a perinuclear region, resembling the Bb in the oocyte, which forms around the centrosome^52^. To determine if the Xvelo aggregates form around the centrosome, we co-expressed Xvelo with centrosome proteins, mCherry-DZIP1^67^ and RFP-HYLS^76^ (SFig1F). Indeed, we found that the Xvelo assembled around the centrosome in HEK293T cells. Previous Fluorescence recovery after photobleaching (FRAP) experiments have demonstrated that in *Xenopus* and zebrafish, Xvelo/Buc protein forms a stable matrix for the Bb^39,50^. We thus performed a FRAP assay to test if these Xvelo assemblies had similar properties. We found that these Xvelo aggregates, like the Xvelo in oocyte Bb, recovered poorly after photobleaching (SFig1G).

Inspired by the above findings, we further characterized this ‘Bb-like’ structure. It is well-known that Bb accumulates organelles such as mitochondria and ER, and sequesters the germ plasm components^22,24,26^. In fact, the Bb was occasionally referred to as the mitochondrial cloud (MC) in *Xenopus*^77^. We thus tested if the Xvelo assemblies in HEK293T cells had the ability to recruit mitochondria, ER, and germ plasm components. In un-transfected cells, the mitochondria and ER are spread throughout the cell. In the cells with Xvelo assembly, however, the mitochondria and ER are selectively enriched into the Xvelo assembly (Fig 1A-B). Thus, overexpression of the Xvelo protein alone was sufficient for re-constituting a ‘Bb-like’ structure in HEK293T cells. In order to further test if this function of Xvelo is specific to the context of HEK293T cells, we performed similar experiments in HeLa cells. We observed that Xvelo assembled around the centrosome and enriched mitochondria and ER in HeLa cells as well (SFig2A-C). Thus, the ability of Xvelo assembly to sequester organelles does not require any oocyte-specific factors and is independent of cell context.

**Figure 1:**
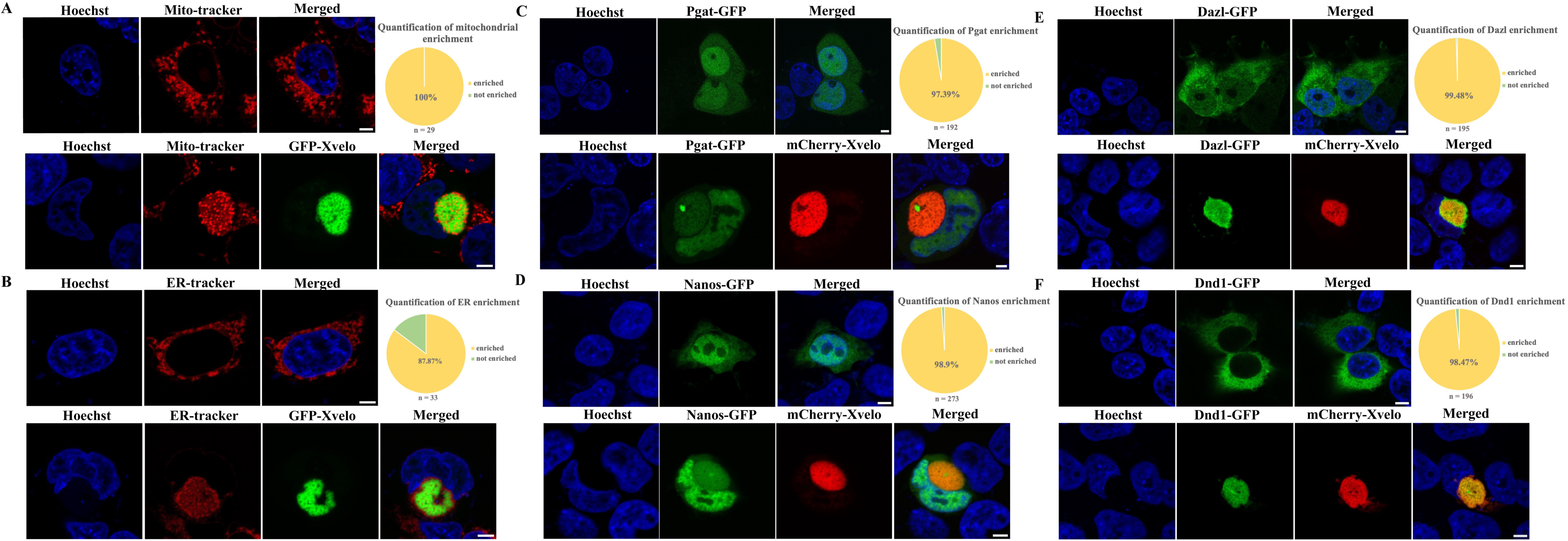
GFP-Xvelo assembles into ‘Balbiani body-like’ aggregates in HEK293T cells. **A** and **B** are representative images to show the subcellular distribution of mitochondria (**A**) and ER (**B**) in control cells (upper panels) and GFP-Xvelo transfected cells (lower panels). Both mitochondria and ER were spread in the entire cytoplasm in control cells, but were recruited into the GFP-Xvelo assembly in GFP-Xvelo transfected cells. The pie graphs at the upper right corner are the quantification of recruitment into the assembly. **C** - **F** are representative images to show the subcellular distribution of germline determinants, including Pgat (**C**), Nanos (**D**), Dazl (**E**), and Dnd1 (**F**). Upper panels show the distribution of germline determinants when they were expressed alone in HEK293T cells. Lower panels show their distribution when co-expressed with GFP-Xvelo. The pie graphs at the upper right corner are the quantification of assemblies in which germline determinants colocalize with Xvelo. Scale bars: 5µm.

One of the most important roles of the Bb is to enrich germline determinants and control their asymmetric localization inside the oocyte. To determine if the Xvelo assemblies in somatic cells functionally behave like the Bb, we asked if they could selectively recruit germ plasm components. To test this, we transfected cells with Xvelo, along with germ plasm components, including Pgat^78^, xDazl^79^, Nanos1^80^, or Dnd1^81,82^. When expressed alone in HEK293T cells, Pgat and Nanos1 were distributed in both the nucleus and cytoplasm, with a higher level of protein being detected in the nucleus (Fig1C and D). When co-expressed with Xvelo, all cytosolic Pgat and Nanos1 proteins were enriched in the Xvelo assembly (Fig1C and D). Different from Pgat and Nanos1, xDazl was highly enriched in the cytoplasm (Fig 1E), and Dnd1 was exclusively present in the cytoplasm (Fig 1F). In cells with Xvelo assembly, xDazl and Dnd1 were completely enriched in the Xvelo assembly (Fig 1E and F). We counted the percentage of cells in which these germ plasm components colocalized with Xvelo aggregates. We found that 97.4% (n=192) of Pgat, 98.9% (n=273) of Nanos1, 99.5% (n=195) of xDazl, and 98.5% (n=196) of Dnd1 were enriched in Xvelo aggregates (Fig 1C-F). Additionally, this ability of Xvelo assemblies to enrich germ plasm components is not specific to HEK293T cells. We observed colocalization of xDazl with the Xvelo aggregates in HeLa cells (SFig2E). To determine if Xvelo assemblies can recruit non-germline proteins, we co-expressed GFP-myc and hRusc-GFP^83^ with mCherry-Xvelo. Neither GFP nor hRusc-GFP colocalized with the Xvelo aggregates (SFig3). Based on the above findings, we conclude that Xvelo can function as an organizer to assemble a “functional” Bb-like structure in somatic cells.

### Centrosome proteins promote assembly of Xvelo

Even after more than a century of the identification of the Bb^21,84^, we do not fully understand the mechanism of its initial assembly. From ultra-structural studies, it was noted that the Bb assembles near the nucleus, around the centrosome^21,51,85^. A recent study showed that dynein-mediated transport of Buc condensates along the microtubules regulates the growth and maturation of Bb in zebrafish oocytes^50^. In our studies in cell culture, we noticed that the Xvelo assembles around the centrosome. This led us to speculate that centrosome components could potentially assist in Bb assembly. Since the Xvelo assemblies in somatic cells resemble the endogenous *Xenopus* and zebrafish Bb, we took advantage of this system to determine if centrosome components could assist in the assembly of Bb. We took a candidate approach and selected a few proteins that localize to different parts of the centrosome. We co-expressed these proteins in cells with Xvelo and tested if any of these components could influence the assembly of Xvelo by quantifying the percentage of cells with aggregated or smeared Xvelo (Fig 2A). Xvelo, on its own, formed aggregates in 10% or fewer cells. We found that overexpression of Sas6^65^, Cenexin^86,87^, DZIP1^67^, or Polo-like kinase 4 (PLK4)^60,88^ were able to promote assembly of Xvelo. Sas6 showed the most prominent effect with nearly 40% of the cells forming aggregates. Another centrosome protein Cep164^89^, however, showed minimal effect on Xvelo assembly (Fig 2A). We further validated these results biochemically by comparing insoluble and soluble Xvelo protein in a fractionation assay (same as SFig1C). Overexpression of Sas6, DZIP1, and Cenexin increased Xvelo in the insoluble (pellet) fraction (Fig2B and C). These results suggest that a subset of centrosome proteins could promote Xvelo assembly around the centrosome.

**Figure 2:**
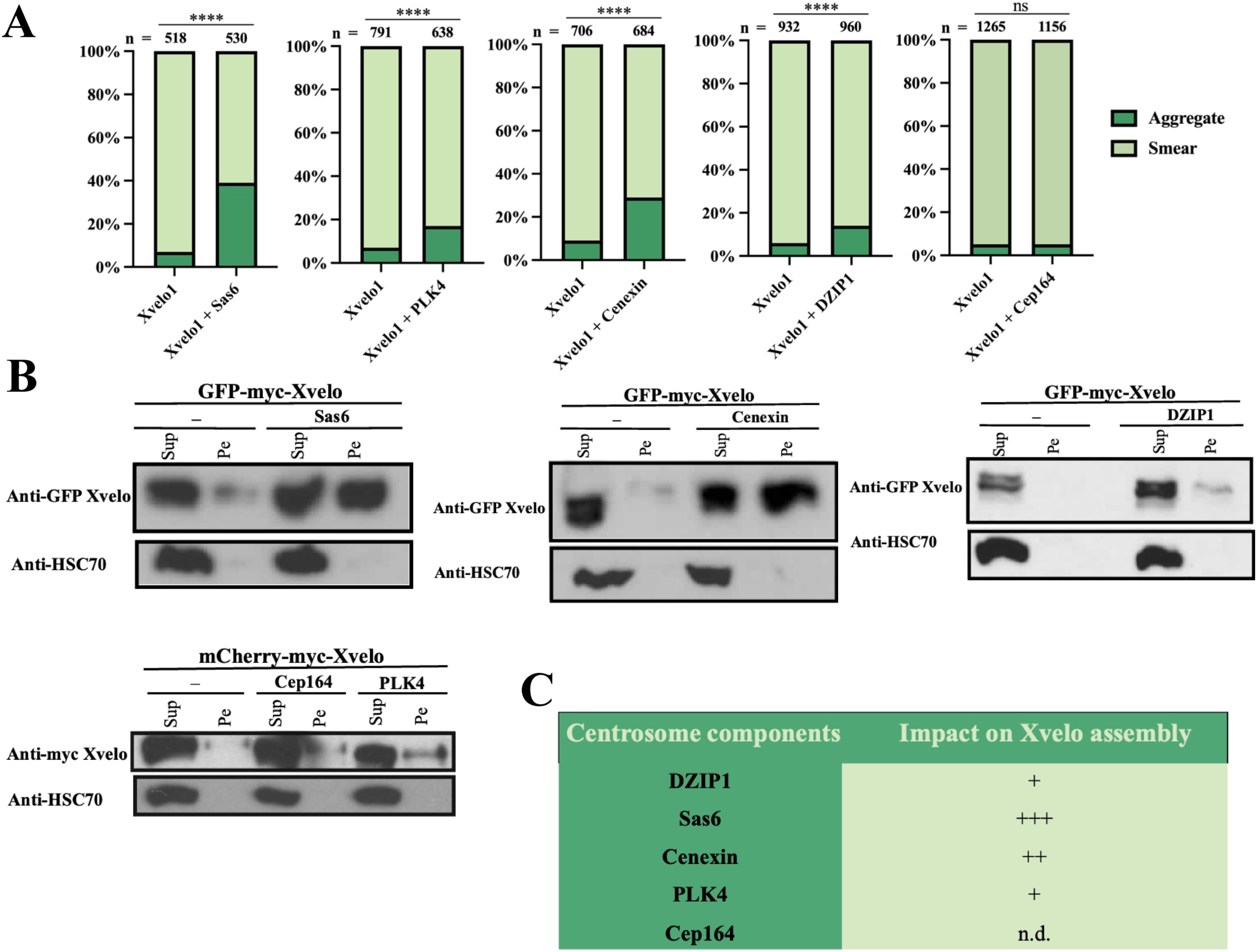
Co-expression of centrosome components can promote the assembly of Xvelo. **A.** Bar graphs show the effects of overexpression of Sas6, PLK4, Cenexin, DZIP1, and Cep164 on the aggregation of Xvelo. The numbers of cells counted are listed on top of each bar graph. Fisher’s exact test was performed for statistical analysis (****p<0.0001). **B.** The effects of overexpression of Sas6, PLK4, Cenexin, xDZIP1, and Cep164 on the assembly of Xvelo were assessed by fractionation and Western blot. HSC70, a protein highly enriched in the soluble fraction, was used as a loading control. **C.** Summary of the impact of different centrosome components on Xvelo assembly. n.d. - not detected.

### Centrosome proteins act on different domains of Xvelo to promote its assembly

Since Sas6, DZIP1 and Cenexin promote the assembly of Xvelo, we explored their interaction with Xvelo. Xvelo has a PLD in the N-terminus, which is involved in its self-assembly, and a C-terminal IDR^39^ (Fig 3A). We started by testing if the N-(termed as F12) or the C-terminal (F34) half of the protein was sensitive to the over-expression of Sas6, DZIP1, and Cenexin. We found that over-expression of Sas6 and Cenexin promoted the assembly of both F12 and F34, while DZIP1 could only induce assembly of F34 (Fig 3B). Notably, the effect of Sas6 on the assembly of F12 was much stronger than that of Cenexin. To determine if Sas6 and Cenexin act on the PLD to induce Xvelo assembly, we performed a similar experiment with F1, which contains the PLD, and F2, which lacks the PLD (Fig 3A). Indeed, we found Sas6 and Cenexin increased the amount of insoluble F1 without affecting the solubility of F2 (Fig 3C). Thus, Sas6 and Cenexin can act on the PLD to induce Xvelo assembly. Since Sas6, DZIP1, and Cenexin can promote the assembly of the C-terminal of Xvelo protein, we further tested their effects on the assembly of F3 and F4 fragments. We found that overexpression of Sas6, DZIP1, and Cenexin could increase the amount of insoluble F3 and F4 (Fig 3C). As Sas6, Cenexin, and DZIP1 can act on different domains of Xvelo to induce Xvelo assembly, we went on to determine if these centrosome proteins physically interact with Xvelo. We generated GST-tagged Xvelo F1, F3, and F4 and performed GST-pull down experiments. We found that Sas6 interacted with F1, F3, and F4 of Xvelo, whereas Cenexin and DZIP1 interacted with F3 and F4 (Fig 3D). Collectively, our results show that different proteins in the centrosome can interact with Xvelo and promote its assembly. We speculate that by binding to and enhancing Xvelo assembly, centrosome proteins assist in the formation of Bb around the centrosome early in oogenesis.

**Figure 3:**
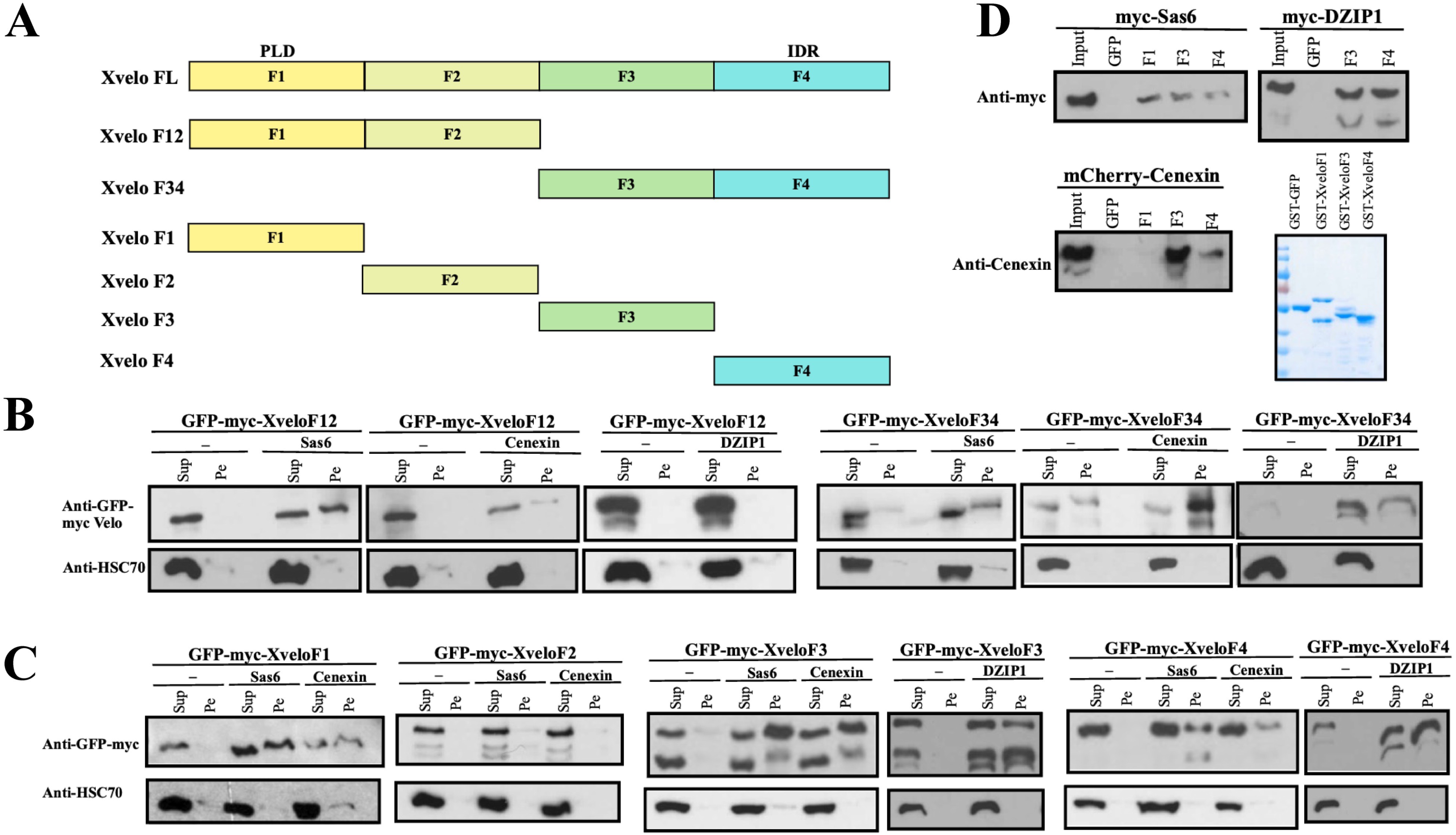
Centrosome proteins act on different domains of Xvelo to promote its assembly. **A.** Schematic representation of the full length Xvelo (779 amino acid residues) and different fragments are shown. Xvelo F12 (1-410 aa), Xvelo F34 (386-779 aa), Xvelo F1 (1-160 aa), Xvelo F2 (138-410 aa), Xvelo F3 (386-610 aa) and Xvelo F4 (586-779 aa). **B.** The effects of overexpression of Sas6, Cenexin, PLK4, DZIP1, and Cep164 on the aggregation of N terminus (F12) or C terminus of Xvelo (F34) were assessed by fractionation and Western blot. **C.** The effects of overexpression of Sas6, Cenexin, PLK4, DZIP1, and Cep164 on the assembly of the different fragments of Velo (F1, F2, F3 and F4) Xvelo were assessed by fractionation and Western blot. **D.** GST pull down and Western blot was performed to test for physical interaction between Xvelo fragments and Sas6, Cenexin and DZIP1. Coomassie brilliant blue staining to estimate protein concentration is shown in the lower right panel.

### Knockdown of centrosome proteins reduces Xvelo assembly

The above studies demonstrate that Xvelo assembles into a Bb-like structure around the centrosome and a subset of centrosome proteins, including Sas6, DZIP1, and Cenexin, can interact with and promote Xvelo assembly. We further determine if Sas6, DZIP1, and Cenexin are required for Xvelo assembly in HEK293T cells. We performed shRNA-mediated knockdown of Sas6, DZIP1, and Cenexin (SFig4) and asked if depleting these centrosome proteins can deter the assembly of Xvelo. For this experiment, HEK293T cells were transfected with GFP-Flag-Xvelo and then infected with lentiviral shRNAs against *sas6*, *dzip1*, and *cenexin.* Three days post-infection, cells were harvested for fractionation and western blot analysis (Fig 4A). We found that while the knockdown of DZIP1 only had a marginal effect, the knockdown of Sas6 and Cenexin dramatically reduced the amount of Xvelo in the insoluble fraction (Fig 4B). Knockdown of Sas6, Cenexin, and DZIP1 in combination was able to reduce the amount of Xvelo in the pellet even further (Fig 4B). The fact that Sas6, Cenexin, and DZIP1 are required for Xvelo assembly provides strong support for our hypothesis that the centrosome assists in Bb formation by promoting Xvelo assembly.

**Figure 4:**
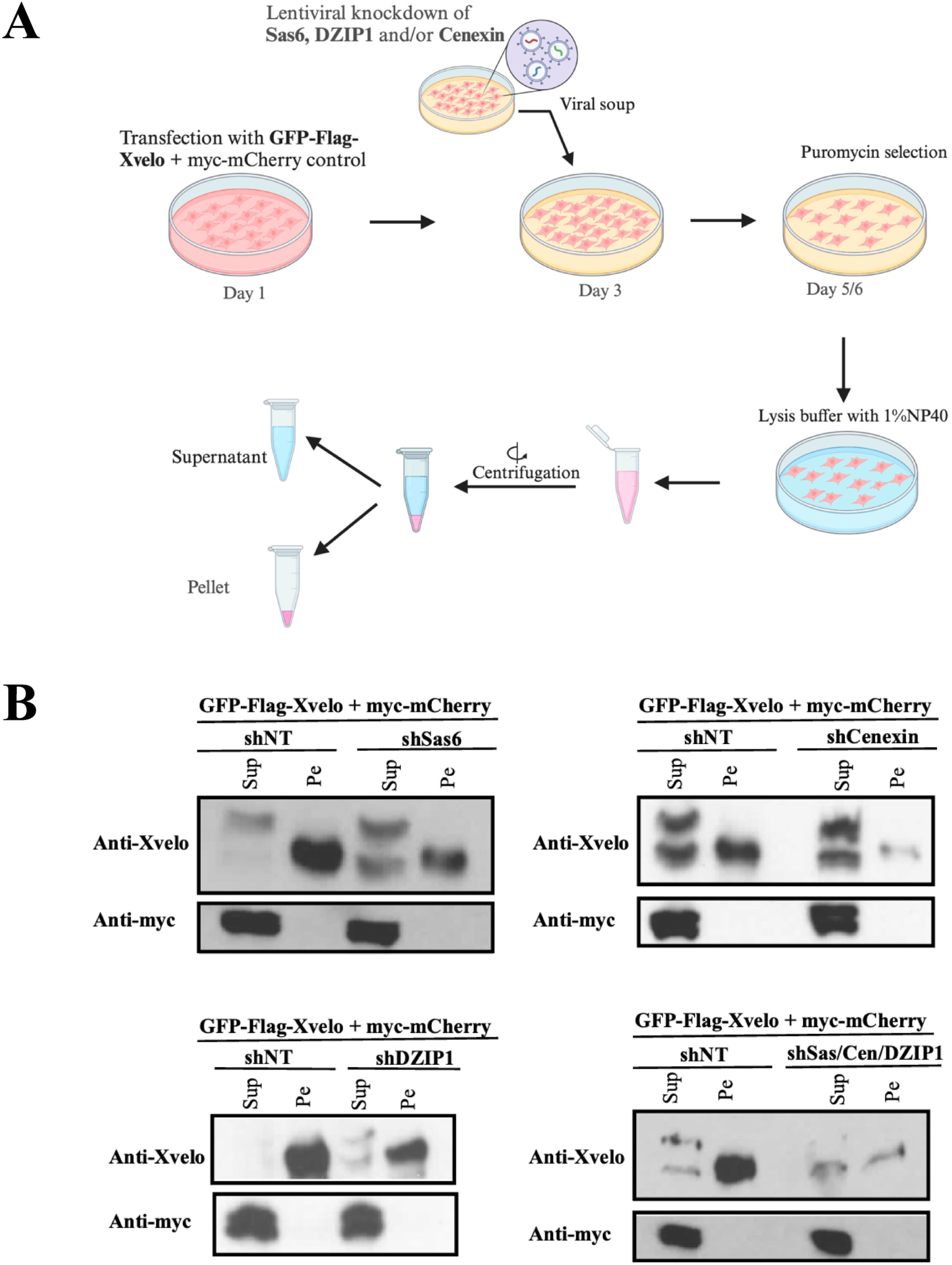
Knockdown of centrosome components impairs the ability of Xvelo to assemble. **A.** Schematic for the experimental plan of knockdown experiment. **B.** Knockdown of Sas6, Cenexin and DZIP1 individually, or in combination were followed by Fractionation and Western blot. Western blots to assess the assembly of Xvelo are shown. myc-mCherry was used as a control for transfection and loading.

## Discussion

The Bb is a highly conserved physiological amyloid in the oocyte. While the Bb was first discovered in the 1800s^21^, the mechanism governing its initial assembly is not well understood. It is known that the Bb accumulates germline determinants needed for the germ cell specification ^30,33,90^ and is crucial for the proper localization of these determinants in the oocytes of organisms like *Xenopus* and zebrafish^10,22,24^. In these species, the initial assembly of the Bb occurs around the centrosome during early oogenesis^51,52^. This event also marks the point of oocyte symmetry break and the establishment of the animal/vegetal axis of the oocyte^30,33,44^. Subsequently, the Bb breaks into numerous small ‘germ plasm islands’, which are transported to and stored at the vegetal cortex of the oocyte^90^. During the oocyte-to-embryo transition, vegetally localized germ plasm undergoes drastic remodeling. Ultimately, the majority of ‘germ plasm islands’ are eliminated in the somatic tissues^91,92^. The remaining islands coalesce into large aggregates and are inherited by a few cells in the embryo, leading to the specification of the primordial germ cells. It is known that *Xenopus* Xvelo and its zebrafish homolog Buc function as the matrix protein for the Bb and germ plasm^33,39,49^. In fact, when overexpressed in fertilized eggs where all germ plasm components needed to specify primordial germ cells were available, Buc was sufficient for germ plasm assembly, inducing ectopic PGCs in early zebrafish embryos ^33,40,41^. Nonetheless, it was unclear if Buc alone is sufficient for the assembly of a Bb in the absence of other germline components.

In this work, we study the assembly of Xvelo in mammalian somatic cells, which lack germ plasm components and other oocyte-specific factors. We found that Xvelo alone was sufficient to reconstitute a ‘Bb-like’ structure in the mammalian somatic cells. These Bb-like assemblies are reminiscent of the Bb in *Xenopus* and zebrafish early oocytes. They form around the centrosome and are capable of recruiting cellular organelles such as mitochondria and ER. Moreover, these Xvelo assemblies can enrich co-expressed germ plasm components such as Dazl, Pgat, Dnd1, and Nanos without affecting the subcellular distribution of non-germline factors. Our FRAP experiment further demonstrates that these Bb-like structures were extremely stable and recovered poorly after photo-bleaching. These findings strongly argue that the assembly of a Bb relies entirely on the biophysical properties of Xvelo/Buc.

The Xvelo protein has a N-terminal PLD that can self-assemble into micron-scale amyloid-like networks in vitro^39^. In *Xenopus* oocytes, the PLD is necessary and sufficient for the localization of Xvelo to the Bb. Interestingly, the PLD alone cannot recruit mitochondria and mRNAs. The C-terminal region of Xvelo is essential for these functions^39^. Our work here reveals that both the PLD and the C-terminal portion of Xvelo are required for the assembly of a Bb-like structure in mammalian somatic cells. The PLD can promote self-assembly while interacting with centrosome components like Sas6 to further increase Xvelo assembly. The IDR-bearing C-terminal region of Xvelo can provide interaction opportunities with additional centrosome components like DZIP1 and Cenexin. We speculate that the interactions between Xvelo and these centrosome proteins are important, as they can recruit Xvelo to the centrosome, initiate Xvelo assembly, and thus, initiate Bb formation. In support of this view, we found that overexpression of Cenexin, Sas6, and DZIP1 enhanced Xvelo assembly. Furthermore, the knockdown of Cenexin, Sas6, and DZIP1 decreased the assembly of Xvelo. As multiple centrosome components can interact with Xvelo and promote its assembly, the centrosome can act as a Xvelo assembly center to direct the initiation of the Bb formation.

During the final stage of our manuscript preparation, Kar et al^50^ reported a mechanism for the maturation of the Bb in zebrafish. It was demonstrated elegantly that once Bb assembly is initiated, small Buc protein condensates in the oocyte utilize a dynein-mediated transport mechanism to join the existing Bb, leading to the growth and maturation of the Bb. Thus, our findings presented here and the results in the literature collectively support the following model. During early stages, centrosome proteins such as Sas6, Cenexin, and DZIP1 promote assembly of Xvelo/Buc to initiate Bb formation around the centrosome. Subsequently, the centrosome acts as MTOC to facilitate dynein-dependent transport of Buc/Xvelo granules^50^. This eventually gathers all Xvelo/Buc protein condensates around the centrosome, leading to the formation of a mature Bb in the perinuclear region of the oocyte.

It is worth mentioning that our model is based on experiments conducted in mammalian somatic cells. Due to the scope of the current work, we only knocked down Cenexin, Sas6, and DZIP1 and investigated their roles in Xvelo assembly in HEK293T cells. In the future, we will carry out loss-of-function experiments in the oocyte and determine if the centrosome facilitates the initiation of Bb formation through the interaction between centrosome proteins and Xvelo/Buc. In addition, since Xvelo interacts with Sas6, Cenexin and DZIP1, and possibly some other centrosome proteins, it will be critically important to investigate the dynamic interactions between centrosome and Xvelo/Buc using high-resolution imaging approaches. Despite the fact that further work is clearly needed, our study presented here provides a novel model to explain the molecular mechanism governing the initiation of Bb assembly.

## Materials and Methods

### Methods

#### Cell culture and transfection

HEK293T cells and HeLa cells were maintained in Dulbecco’s Modified Eagle Media (DMEM) supplemented with 10% Fetal Bovine Serum (FBS) and 50IU/ml penicillin-streptomycin. For transfections, DMEM was replaced with Opti-MEM™ and transfections were carried out using Polyethylenimine (PEI).

#### Cellular fractionation

HEK293T cells were transfected with Xvelo (0.5µg) in addition to either a control plasmid or centrosome components (0.5µg). 24 hours post-transfection, the cells were washed with cold PBS, followed by lysis on ice using 1% NP40 lysis buffer (50mM Tris pH 7.6, 125mM NaCl, 1mM EDTA, 1% NP40%, and Protease inhibitor cocktail) for 10 minutes. Collected lysates were subjected to centrifugation at 13,000 rpm for 5 minutes on a tabletop centrifuge, followed by separation of supernatant and pellet fractions. Samples were further processed with SDS Sample buffer for western blot.

#### Western Blot

Western blot samples were boiled for 5 minutes at 95°C, left on ice for two minutes, followed by a quick spin. The supernatants were loaded on SDS-PAGE gel. After the electrophoresis, proteins were transferred to a PVDF membrane. For blocking, PVDF membranes were incubated with 5% nonfat dry milk (in 1XTBST: Tris-buffer saline + 0.1% Tween 20). This was followed by overnight incubation with primary antibody, and secondary antibody incubation the next day. After washing with 1XTBST extensively, the membranes were developed using ECL™ Prime Western Blotting Detection Reagent.

#### Mito-tracker and ER tracker staining

HEK293T or HeLa cells were seeded on glass bottom dish and transfected the next day with 0.5µg Xvelo. One day post-transfection, the cells were incubated in Fluorobrite™ DMEM with 100nM mito-tracker or 1000nM ER tracker for 30min, along with Hoechst to visualize the nuclei. Xvelo aggregates and mitochondria or ER were then imaged using the Nikon A1R confocal microscope.

#### Fluorescence Recovery After Photobleaching (FRAP)

HEK293T cells were seeded on glass bottom dish and transfected the next day with 0.5µg Xvelo. One day post-transfection, DMEM was replaced with pre-warmed Fluorobrite™ DMEM and Nikon A1R confocal microscope was utilized to perform the FRAP experiment. An ROI was selected on the Xvelo aggregates that were subject to photobleaching using lasers with 50% power. The recovery of Fluorescence after bleaching was monitored every 30 seconds, for about 10 minutes. ImageJ was used for further analysis involving measurement of fluorescent intensity over time in the bleached region and normalized with respect to an unbleached region of the same size on the aggregate.

#### Lentiviral Knockdown

HEK293T cells were transfected with viral packaging vectors pAX2 and pMD2, along with a non-target scrambled control shRNA or shRNAs against Sas6, Cenexin (ODF2) and DZIP1. All shRNA vectors were purchased from Sigma Aldrich. 24 hours post-transfection, the viral soup was collected every 12 hours for 3 days and stored at 4°C till the cells were ready to be infected. Meanwhile, some HEK293T cells were transfected with Xvelo and a control plasmid. Two days after transfection, the transfected cells were infected with either a non-target control virus or viral soup for knockdown of centrosome components (Sas6, Cenexin and/or DZIP1). For knockdown of multiple centrosome components, the cells were infected with virus for shSas6, shCenexin, and shDZIP1 sequentially for 24h each. Puromycin selection was allowed overnight using 10µg/ml Puromycin. To validate the knockdown, RNA from cells was extracted using TRIzol reagent. This was followed by cDNA synthesis using M-MLV Reverse Transcriptase enzyme, and real-time PCR using 2X SYBR Green qPCR Master Mix. The Applied Biosystems QuantStudio 3 Real-Time PCR system was utilized to obtain Ct values. The primers for qPCR are listed under Materials (STable1).

#### GST pulldown

pGEX-6p1-GST-F1, pGEX-6p3-GST-F3, pGEX-6p3-GST-F4 and pGEX-6p1-GST-GFP were expressed in BL21 cells, followed by induction with 1mM IPTG for 4h at 30°C. Following sonication, the cleared lysates were incubated with Glutathione Sepharose beads for 3 hours at 4°C. After thoroughly washing the beads with lysis buffer (300g for 30 sec each wash), we estimated the amount of protein by Coomassie Brilliant Blue staining. For the pull-down experiments, cleared lysates were obtained from HEK293T cells that were previously transfected with Sas6, xDZIP1, or Cenexin. After saving the ‘Input’, the rest of the lysates were incubated overnight at 4°C with roughly similar amounts of GFP/F1/F3/F4 protein-coated beads. The next day, the beads were washed 4 times with lysis buffer (300g for 30 sec each), 1X SDS Sample buffer was added to the beads, and samples were processed for western blot analysis.

## Materials

### See Key Resource Table (KRT)

**STable1:**
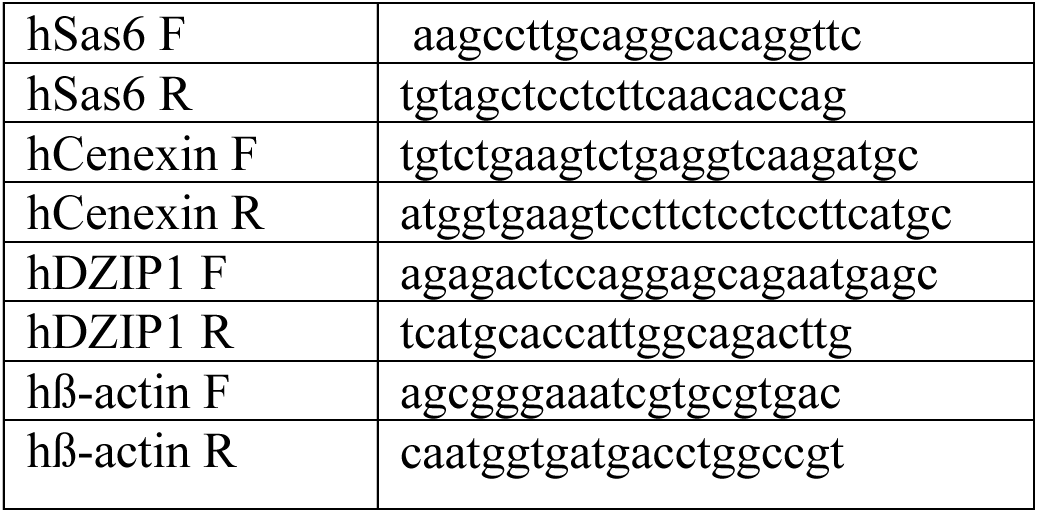
List of oligonucleotides used for knockdown validation.

## Supporting information

SFigure 1

SFigure 2

SFigure 3

SFigure 4

## Acknowledgments

This work is supported by a grant from NIH (R35 GM131810). We appreciate Dr. Brian Mitchell for providing HYLS and Sas6 expression constructs.

**SFigure1: GFP-Xvelo can form assemblies around the centrosome in cultured HEK293T cells.**

**A.** Representative images of Xvelo aggregates and smears in HEK293T cells are shown. **B.** Bar graph shows the quantification of dose-dependent assembly of Xvelo as percentage of cells with smears and aggregates of Xvelo. The numbers of cells counted are listed on top of each bar graph. **C.** Schematic for Fractionation protocol to separate soluble protein and insoluble protein aggregates is shown in the top panel. The effect of increasing Xvelo concentration on the assembly of Xvelo was assessed by fractionation and Western blot in the bottom panel. **D.** Fractionation and Western blot results show gradual accumulation of Xvelo assemblies within the first three days post-transfection. By five days post-transfection, Xvelo assemblies are partially degraded. **E.** Representative images for the effect of digitonin (0.3%) extraction on Xvelo assemblies and GFP control are shown. The cells were treated with 0.3% Digitonin containing PBS for 1 minute. **F.** Co-expression of DZIP1 or HYLS along with GFP-Xvelo was conducted to observe location of Centrosome with respect to Xvelo assembly. **G.** Normalized intensity of GFP fluorescence in the Xvelo aggregates was followed post-photobleaching (n = 6). Representative images are shown in the upper panel. Scale bars: 5µm.

**SFigure2: GFP-Xvelo expression in HeLa cells can induce the formation of ‘Balbiani body-like’ structures of Xvelo.**

**A.** Co-expression of DZIP1 along with GFP-Xvelo was conducted to observe location of Centrosome with respect to Xvelo assembly in HeLa cells. **B** and **C** are representative images to show the subcellular distribution of mitochondria (**B**) and ER (**C**) in control cells (upper panels) and GFP-Xvelo transfected cells (lower panels). Both mitochondria and ER were spread in the entire cytoplasm in control HeLa cells, but were recruited into the GFP-Xvelo assembly in GFP-Xvelo transfected cells. **D-E** are representative images to show the subcellular distribution of non-germline determinants, myc-GFP **(D)** and germline determinant, Dazl **(E)**. Upper panels show the distribution when they were expressed alone in HeLa cells. Lower panels show their distribution when co-expressed with GFP-Xvelo. The pie graph at the upper right corner **(E)** is the quantification of cells in which germline determinant Dazl colocalized with Xvelo assembly. Scale bars: 5µm.

**SFigure3: Xvelo assemblies in HEK293T cells do not selectively enrich non-germline determinants.**

**A-B** Representative images to show the subcellular distribution of non-germline determinants, myc-GFP **(A)** and Rusc2-GFP **(B)** when co-expressed with mCherry-Xvelo protein. The pie graphs at the right corner are the quantification of assemblies in which non-germline determinants colocalize with Xvelo. Scale bars: 5µm.

**SFigure4: Validation of knockdown of Centrosome components**

qPCR validation of knockdown for individual centrosome components (Sas6, Cenexin, DZIP1) in the top panel and knockdown of all three in the bottom panel are shown.

## Key resources table

| REAGENT or RESOURCE | SOURCE | IDENTIFIER |
| --- | --- | --- |
| **Antibodies** |  |  |
| Rabbit anti-Xvelo | Boke et al.<sup>39</sup> | N/A |
| Mouse anti-HSC70 | Santa Cruz | Cat. Sc-7298;<br>RRID:<br>AB_627761 |
| Rabbit anti-mDZIP1 | Jin et al.<sup>73</sup> | N/A |
| Goat anti-GFP | Rockland | Cat. 600-101-215;<br>RRID:<br>AB_218182 |
| Mouse anti-c-Myc | Thermo Fisher | Cat. 13-2500;<br>RRID:AB_25330<br>08 |
| Rabbit anti-ODF2 (Cenexin) | Thermo Fisher | Cat. 12058-1-AP;<br>RRID:AB_21566<br>30 |
| Secondary anti-rabbit-HRP | GE Healthcare | Cat. NA934V |
| Secondary anti-mouse-HRP | GE Healthcare | Cat. NA931V |
| **Chemicals, peptides, and recombinant proteins** |  |  |
| Dulbecco's Modified Eagle Media (DMEM) | Fisher Scientific | Cat. SH30243.01 |
| Foetal Bovine Serum | Fisher Scientific | Cat. FB12999102 |
| Penicillin-Streptomycin | Corning | Cat. 30-001-CI |
| Opti-MEM™ | Thermo Fisher Scientific | Cat. 11058021 |
| Fluorobrite™ DMEM | Thermo Fisher Scientific | Cat. A1896701 |
| Puromycin | Thermo Fisher Scientific | Cat. A1113803 |
| Protease inhibitor cocktail | Sigma | Cat. P8340 |
| Glass bottom dish | MatTek | Cat. P35GC-1.5-14-C |
| Mito-tracker | Thermo Fisher Scientific | Cat. M7512 |
| ER-tracker | Thermo Fisher Scientific | Cat. E34250 |
| Hoechst 33342 | Invitrogen | Cat. H1399 |
| TRIzol reagent | Thermo Fisher Scientific | Cat. 15596018 |
| 2 × SYBR Green qPCR Master Mix | Bimake | Cat. B21203 |
| M-MLV Reverse Transcriptase | Promega | Cat. M1701 |
| ECL™ Prime Western Blotting Detection Reagent | Amersham | Cat. RPN2236 |
| Glutathione Sepharose™ 4B | Sigma | Cat. GE17-0756-01 |
| HEPES | Thermo Fisher Scientific | Cat. BP310 |
| Triton X-100 | Fisher bioreagents | Cat. BP151 |
| Sodium dodecyl sulfate (SDS) | Sigma | Cat. L4390 |
| **Oligonucleotides** |  |  |
| See S-table 1 for a list of oligonucleotides |  |  |
| **Recombinant DNA** |  |  |
| pCS2-GFP-myc-Xvelo | This paper | N/A |
| pCS2-mCherry-myc-Xvelo | Hwang et al.<sup>93</sup> | N/A |
| pCS2-GFP-Flag-Xvelo | This paper |  |
| pCS2-myc-mCherry | This paper | N/A |
| pCS2-RFP-HYLS | Gift from Brian Mitchell | N/A |
| pCS2-RFP-Sas6 | Gift from Brian Mitchell | N/A |
| pCS2-myc-Sas6 | Gift from Brian Mitchell | N/A |
| pCS2-mCherry-Cenexin | Addgene | Plasmid #186435 |
| pEGFP-Cep164 | Addgene | Plasmid #41149 |
| pEGFP-C3-PLK4-3xFLAG | Addgene | Plasmid #69837 |
| pCS2-Dnd1-GFP | Aguero et al.<sup>94</sup> | N/A |
| pCS2-Pgat-GFP | Aguero et al.<sup>94</sup> | N/A |
| pCS2-Nanos-ORF-GFP-SV40 | Aguero et al.<sup>94</sup> | N/A |
| pCS2-Dazl-GFP | Aguero et al.<sup>94</sup> | N/A |
| **Software/algorithms** |  |  |
| ImageJ | Schneider et al.<sup>95</sup> | https://imagej.net/ij/index.html |
| Adobe Illustrator | Adobe Illustrator | https://adobe.com/products/illustrator |
| GraphPad Prism | GraphPad | https://www.graphpad.com/ |
| BioRender | Biorender | https://www.biorender.com |

