## Supplementary figures and images for "Centrosome-assisted assembly of the Balbiani body"

### SFigure 1

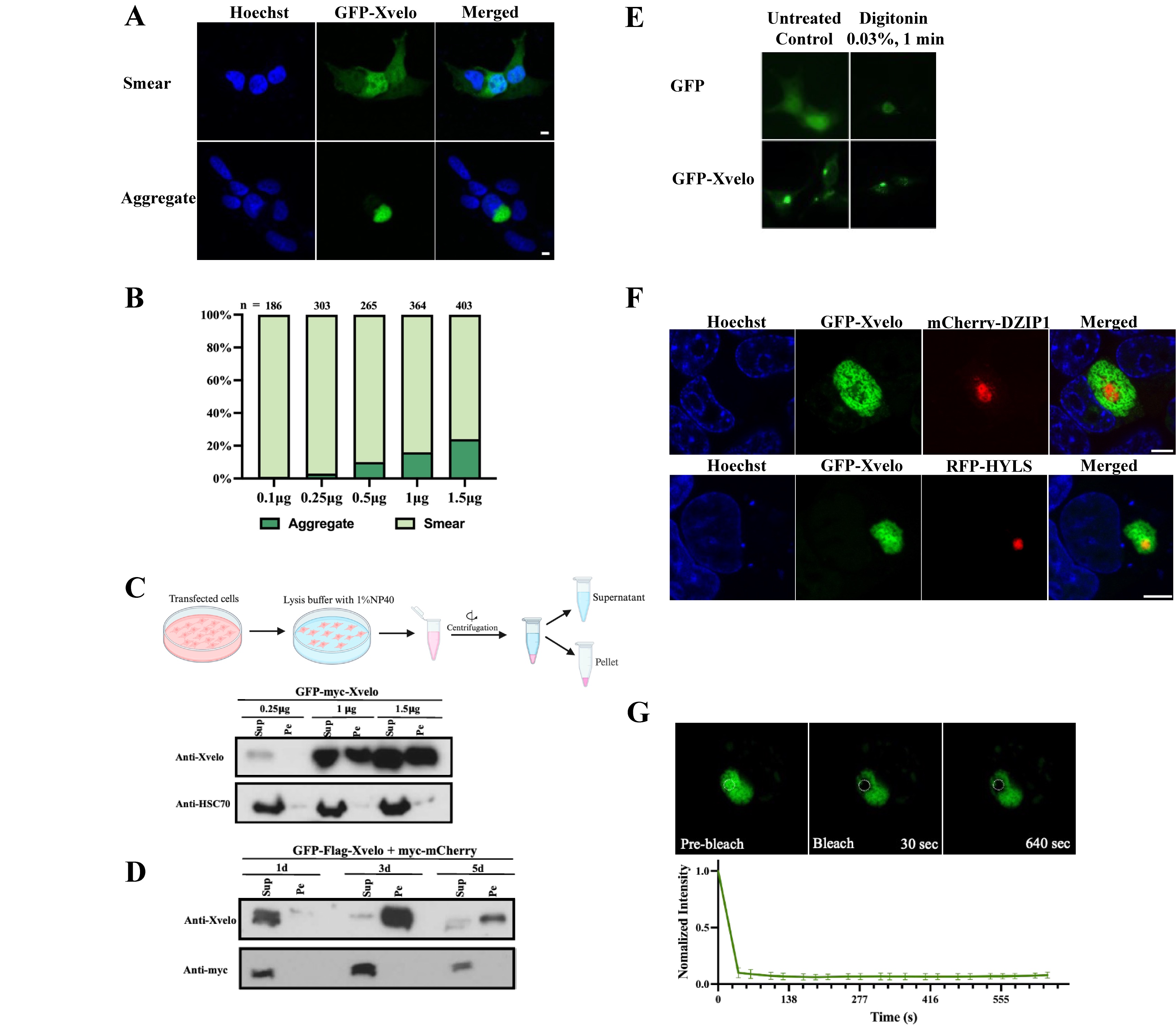

### SFigure 2

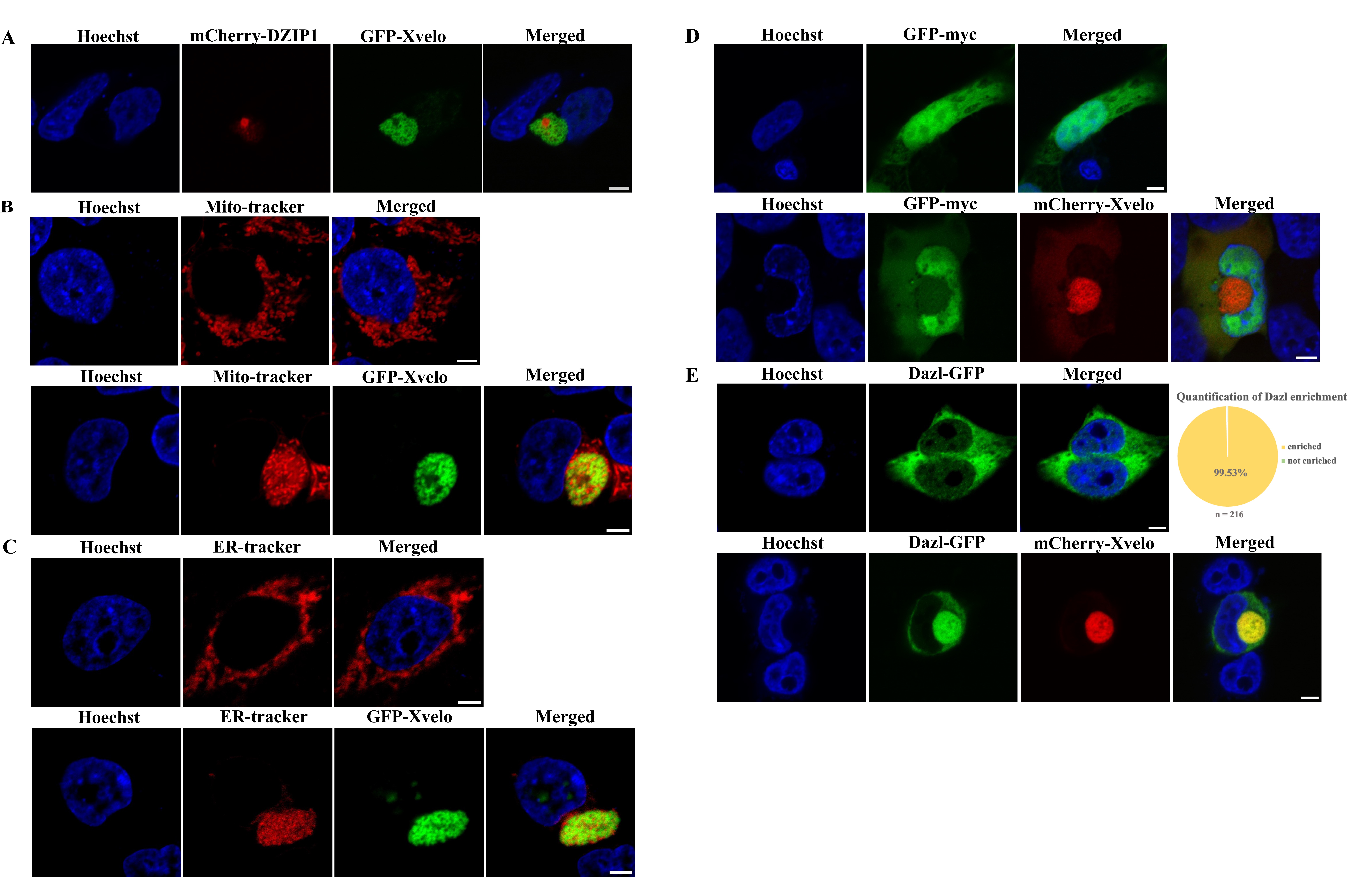

### SFigure 3

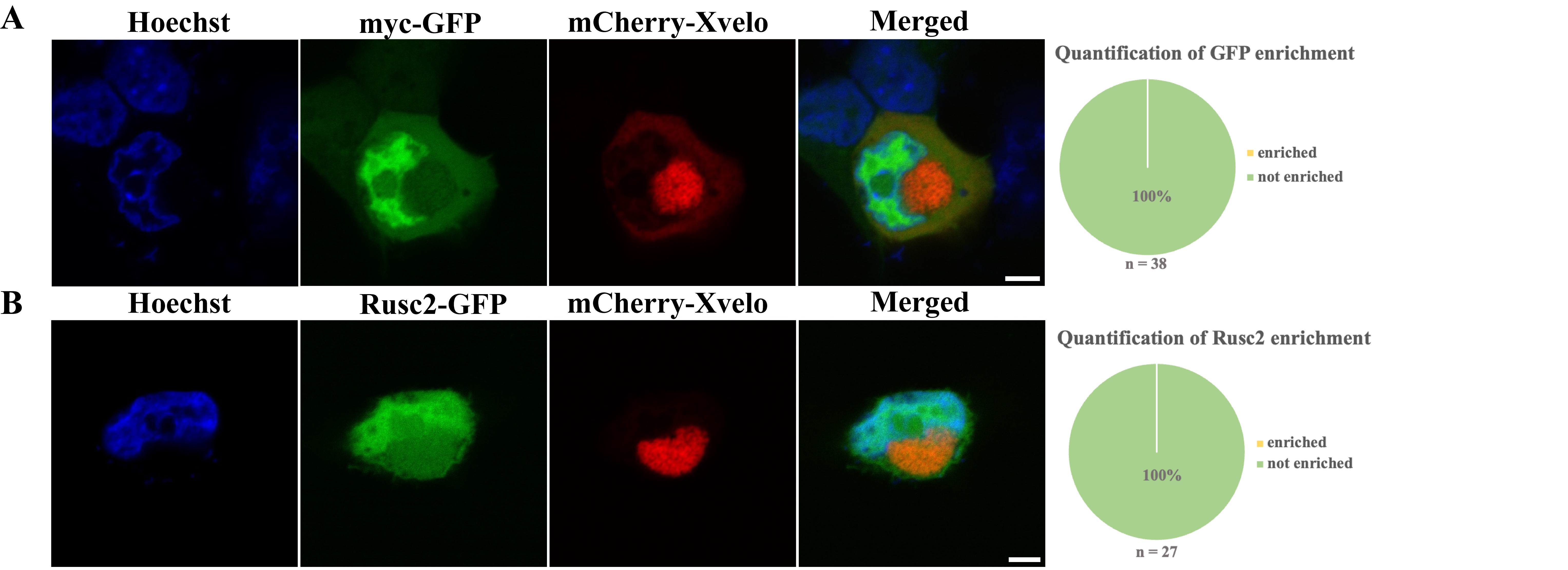

### SFigure 4

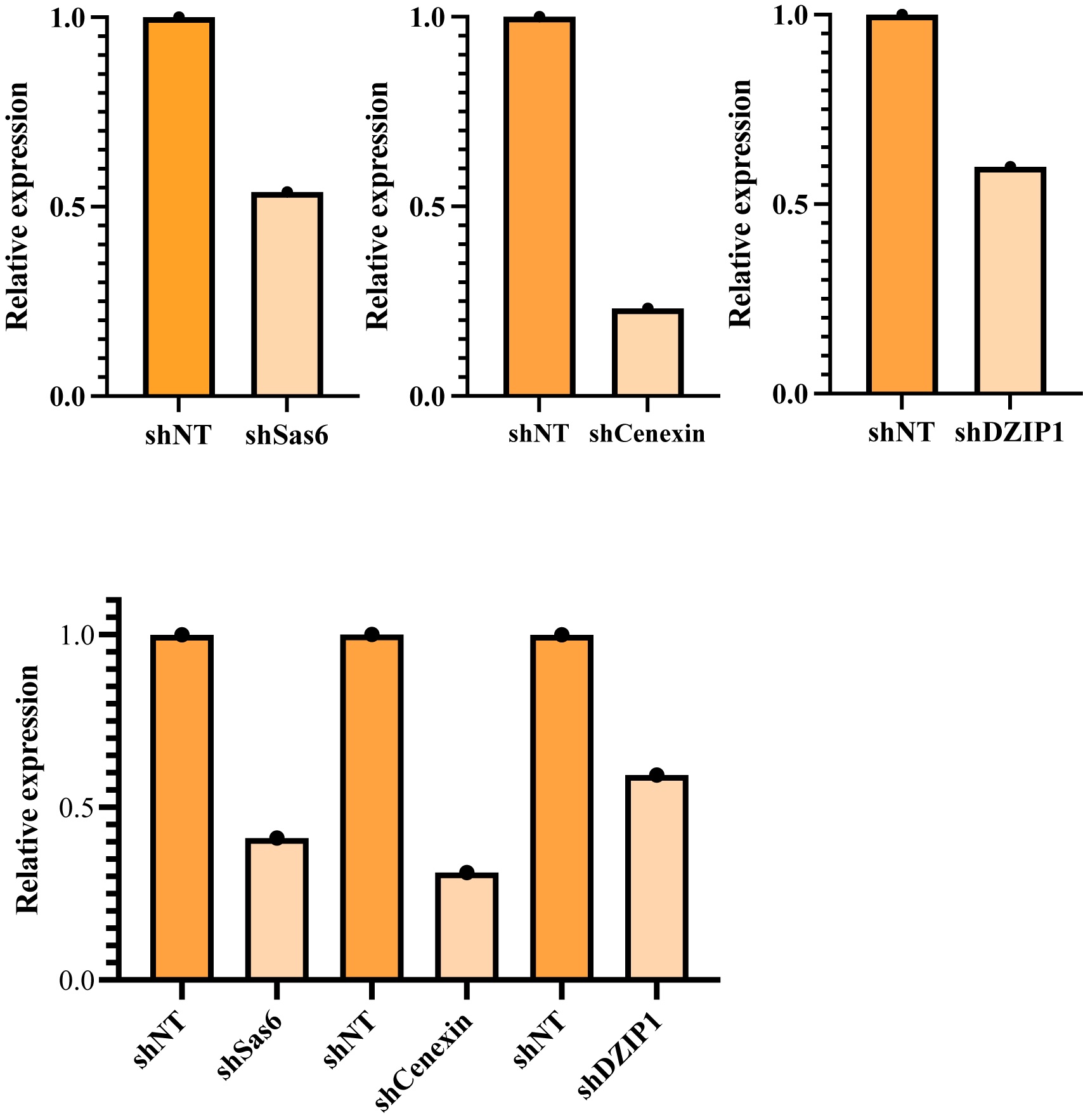
